# Light spectral quality alters glandular trichome architecture more strongly than cannabinoid accumulation in *Cannabis sativa*

**DOI:** 10.64898/2026.06.29.735290

**Authors:** Shaalin Dlaymi, Reilly J. Perovich, Chak-Chung Kuo, Rui Liu, Vincent Fetterley, Alexander Z.-F. Lee, Cory S. Harris, Marco Todesco, A. Lacey Samuels, Marina Cvetkovska

## Abstract

The glandular trichomes in *Cannabis sativa*, found predominantly on female flowers, produce and store a variety of unique phytocannabinoids, increasingly studied for their use in medicinal applications. Maximizing yield and cannabinoid profiles requires the optimization of the environmental factors that regulate plant growth. Light plays a prominent role, both as an energy source but also as an important developmental signal. Thus, optimization of lighting strategies, particularly through customizable light-emitting diode (LED) fixtures, has become a major focus of controlled-environment cannabis research. Here, we focus on the effect of blue-enriched and far red-enriched light spectra on the morphological traits and biochemical profiles of two THCA-dominant varieties: Pineapple Cough and Rocky Fire #7. Spectral composition exerts modest and genotype-specific effects on the plant development, inflorescence biomass, and cannabinoid concentration but we demonstrate a positive correlation between total yield and plant height in both varieties, regardless of spectra. We also show that growth under far-red enriched light affects the visible pigmentation in both varieties with significantly lower chlorophyll levels and paler fan and sugar leaves. Finally, we demonstrate that far-red light consistently increased the trichome stalk length in both varieties, suggesting that spectral composition can alter trichome development and morphology. Our data offers insights into cannabis development and secondary chemical profiles in response to different light spectra, allowing growers to adjust light spectra to obtain desirable cannabis traits for industrial production.

## 1. Introduction

*Cannabis sativa* is an annual herbaceous plant likely native to Central Asia and now cultivated worldwide for fiber, grains, and medicinal applications, as well as cultural, spiritual, and recreational use. The pharmaceutical value of cannabis stems from its ability to synthesize a plethora of unique compounds including cannabinoids, specialized terpenophenolic secondary metabolites (Xie et al., 2023) that interact with the human endocannabinoid system and influence various physiological and psychological responses (Volkow et al., 2017). Cannabinoid accumulation takes place within capitate-stalked glandular trichomes, primarily in pistillate inflorescences and adjacent inflorescence-associated leaves (sugar leaves). The glandular trichomes consist of a multicellular stalk (Siazon et al., 2026) and a glandular head composed of secretory disc cells and an extracellular storage cavity, all covered by a waxy cuticle (Livingston et al., 2020; Mahlberg and Kim, 2004). Cannabinoid biosynthesis occurs within secretory disc cells, where metabolic precursors derived from the methylerythritol phosphate (MEP) and polyketide synthase (PKS) pathways are enzymatically converted to cannabigerolic acid (CBGA) (Gülck and Møller, 2020). Most major cannabinoids, including tetrahydrocannabinolic acid (THCA), cannabidiolic acid (CBDA), and cannabichromenic acid (CBCA), are synthesized from CBGA by cannabinoid synthases in the cell wall between the secretory disk cells and storage cavity, and accumulate in the extracellular storage cavity of the trichome head (Livingston et al., 2020; Sirikantaramas et al., 2005). Therefore, the cannabis inflorescences and the associated glandular trichomes are the most pharmaceutically valuable component of the plant.

Modern cannabis cultivation relies heavily on controlled indoor systems in which environmental factors can be precisely regulated to optimize plant growth and metabolite production. Light shapes nearly all aspects of plant growth and function, with light intensity, photoperiod, and spectral quality each exerting distinct effects on photosynthetic physiology, development, and metabolism (Lauria et al., 2024; Paradiso and Proietti, 2022). Consequently, optimization of lighting strategies, particularly through customizable light-emitting diode (LED) fixtures, has become a major focus of controlled-environment cannabis research aimed at optimizing cannabinoid profiles and yields (Eichhorn Bilodeau et al., 2019; Jin et al., 2019). Cannabis is recognized as a high-light tolerant species that exhibits high photosynthetic capacity, with biomass accumulation increasing across a broad range of light intensities (Chandra et al., 2008; Eaves et al., 2020; Moher et al., 2022).

Flowering, and thus cannabinoid accumulation, in most cannabis varieties is induced by a switch to short photoperiods (Hall et al., 2014; Zhang et al., 2018), allowing growers to control the balance between robust vegetative growth and flowering (Ahrens et al., 2023; Peterswald et al., 2023). While the general relationships between intensity and photoperiod are well established, recent evidence suggest that the interaction between light quality and cannabis growth are more complex than previously appreciated (recently reviewed in Ahsan et al., 2024). Spectral quality, defined as the relative distribution of wavelengths across the visible spectrum, is increasingly recognized as an important regulator of plant morphology, physiology, and secondary metabolism (Huq et al., 2024; Lauria et al., 2024; Wu et al., 2025); however, the mechanisms by which light quality influences development and secondary metabolism in cannabis remain poorly understood.

Most cannabis research to date has focused on the effects of light spectrum on inflorescence yield and cannabinoid accumulation. In many plant species, the ratio of red (600-700 nm) to far-red (700-800 nm) light (R:FR) regulates plant architecture and flowering, and reductions in R:FR induce shade avoidance responses such as stem elongation and reduced branching (Ballaré and Pierik, 2017; Devlin, 2016). In cannabis, reduction in the R:FR ratio through far-red light enrichment promoted stem elongation (Peterswald et al., 2025), consistent with the shade-avoidance response in other plant species. Surprisingly, even red-enriched spectra were also reported to produce taller plants, despite higher R:FR ratios (Danziger and Bernstein, 2021). However, decreasing the R:FR ratio by supplementation with far-red (Kotiranta et al., 2025; Magagnini et al., 2018) or increasing R:FR by supplemental red light (Morello et al., 2022) during growth have been reported to lead to decreased inflorescence biomass, and variable genotype-specific changes to cannabinoid concentrations in different experiments. An interesting approach was to cultivate plants under broad spectrum light with only end-of-day far-red supplementation, which led to genotype-specific cannabinoid concentrations, but ultimately produced no difference in total cannabinoid yield (Peterswald et al., 2025). Thus, there appear to be trade-offs between cannabis plant architecture and yield when supplementing with longer wavelengths although these responses may differ across genotypes.

Shorter wavelengths, particularly blue light (400-500 nm), promote compact morphology and increased photosynthetic capacity through enhanced chlorophyll accumulation and stomatal regulation in many plant species (Fantini and Facella, 2020; Mehmood et al., 2026). In cannabis, blue-enriched spectra have sometimes been associated with more compact plants (Morello et al., 2022), but in a study that systematically increased the fraction of blue photons, significant changes in plant height were not observed (Westmoreland et al., 2021). In some genotypes, blue light induced changes in the chemical make-up of the plant (e.g. increased CBGA but not THCA in drug-type varieties; Danzinger and Bernstein, 2021), but cannabinoid profile changes in response to blue light are often genotype-specific (Phillips et al., 2025) or not statistically significant (Westmoreland et al., 2021). The more compact morphology of plants treated with blue light is often correlated with decreased biomass production (Morello et al., 2022; Wei et al., 2021; Westmoreland et al., 2021). Collectively, cannabis studies that manipulated the blue and red/far-red light suggest a complex interplay between light quality, plant morphology, and secondary metabolism, although these responses are often genotype specific and modest.

From a cannabis cultivation perspective, it is important to establish a strong architectural skeleton during the vegetative plant growth phase to support large inflorescences during the flowering phase. Consequently, optimizing light-induced changes in plant architecture and development may lead to increased yields even if these treatments do not directly affect cannabinoid biosynthesis (Namdar et al., 2019). Therefore, there is a strong need to test cannabis growth using commercially relevant and energy-efficient LED lights that are capable of providing different spectral profiles, with the aim of providing cannabis growers with strategies for getting the best quality crops. In this study, we examined the effects of blue and far-red light supplementation on two industry-relevant and THCA-dominant *C. sativa* varieties and investigated the relationships between plant morphology, trichome architecture, and cannabinoid accumulation. Our findings provide novel insights on the effects of light on cannabis trichome morphology and contribute towards the development of effective lighting strategies for commercial indoor cannabis production.

## 2. Materials and Methods

### 2.1 Plant Cultivation and Experiment Design

Two THCA-dominant *Cannabis sativa* varieties, ‘Pineapple Cough’ (PC) and ‘Rocky Fire #7’ (RF) originally developed by Stinky Brothers (Lumby, BC), were used in all experiments. Clones from female mother plants were provided as 6-week-old rooted cuttings by Green Amber Canada (Kelowna, BC). Each cutting was planted in a 5-gallon pot into a mixture of general-purpose soil (Pro-Mix BX, Premier Tech), coco choir, and black earth soil (5:3:2). Plants were cultivated in 1.5 × 1.5 × 1.8 m grow tents (The Living Room, Urban Grow) housed within an environmentally controlled greenhouse and maintained at 22°C/17°C (±2°C) day/night temperature (University of Ottawa) or a secure lab at 20°C (University of British Columbia). Air circulation within each tent was provided by a forced airflow system with carbon filtration. Leaves and branches within 8 cm of the soil surface were removed after six weeks to improve airflow and light penetration. Irrigation and fertilization were adjusted based on the growth stage, plant requirements, and substrate saturation. Soil pH was monitored 6 cm below the surface of the soil using a pH meter (MS02, Sonkir) and maintained at pH 6-7. Electrical conductivity (EC) was monitored 6 cm below the surface and in the irrigation runoff with an EC/CF/TDS meter (EC-3185, Hilitand Ltd). EC values were maintained at 0.8-1.0 mS m^-1^ during the vegetative phase (Week 0-3 after potting) and 1.0-2.0 mS m^-1^ during the flowering phase (Week 4-10 after potting). Fertilization solution (NutriPlus, Miracle-Gro; 1-1.25 mS m^-1^) was provided every second day and gradually increased from ∼0.5L during vegetative growth (24N-8P-16K) to ∼2.5L during flowering (18N-18P-21K) while maintaining appropriate soil pH and EC.

Light intensity, spectra, and photoperiod were controlled within each tent using LED light fixtures (LED One Lighting Systems; Green Amber Canada). During vegetative growth (Week 0-3), plants were maintained under a long photoperiod (18h/6h light/dark) with a total photosynthetic photon flux density (PPFD) at 700±30 µmol m^-2^ s^-1^. To induce flowering (Week 4), plants were transferred to a short-day regime (12h/12h light dark) and total PPFD was increased to 900±30 µmol m^-2^ s^-1^. Light treatments included: (1) full-spectrum white light (430-730 nm, control), (2) white light with blue-enrichment (430-480 nm), and (3) white light with far-red enrichement (730-780 nm) spectra (Fig. 1A; Table S1). Light measurements were obtained from the upper canopy (0.6 m from light source) using a spectrometer (Model LI-180; Li-COR). To avoid location-specific effects the position of the plants within the tents was periodically randomized. Unless otherwise noted, all experiments were done with three biological replicates, each with at least three individual plants as technical replicates.

**Figure 1.**
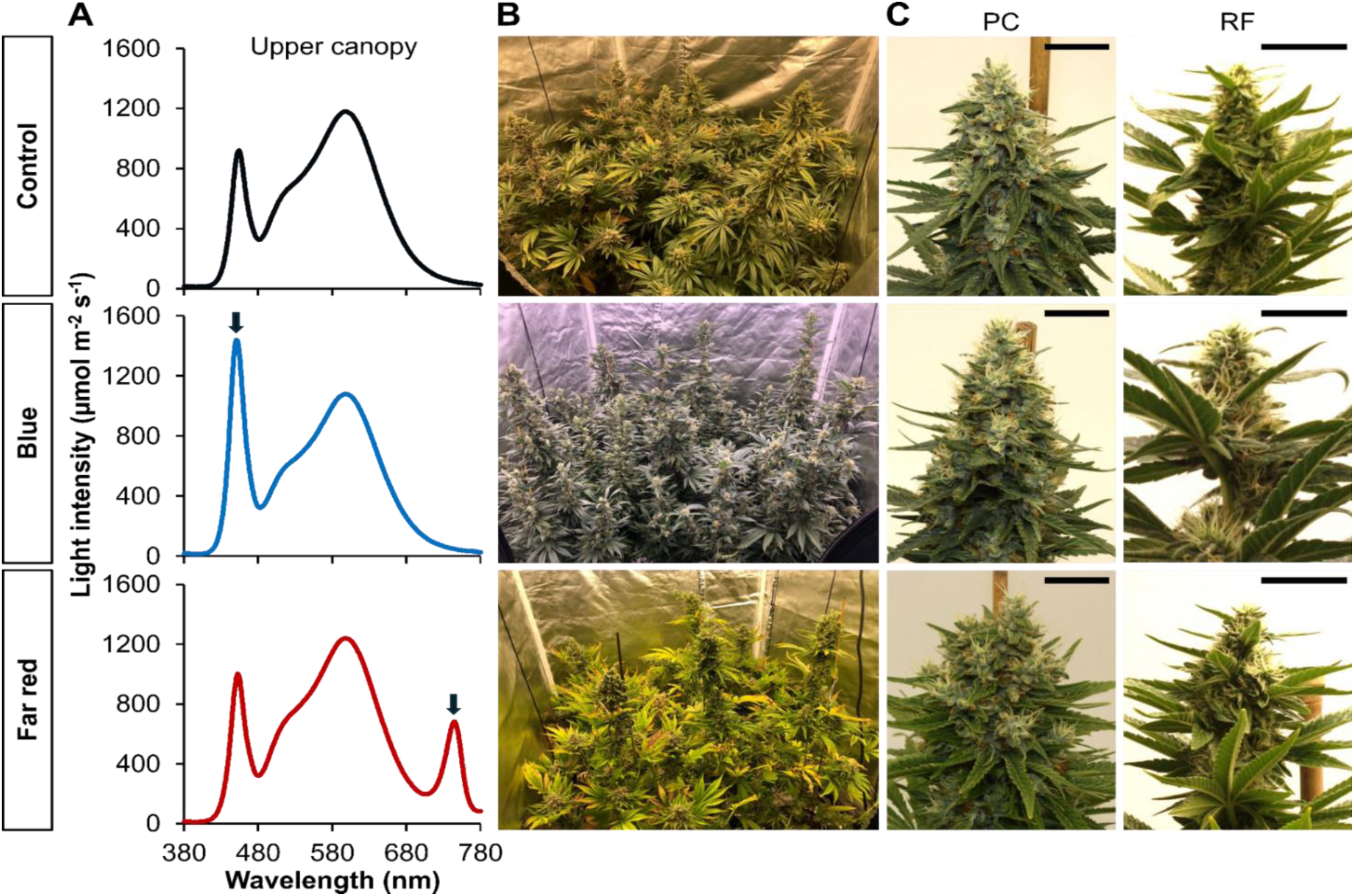
Light spectra and cannabis plant morphology. **(A)** Light spectra for the control, blue-enriched, and far-red-enriched treatments measured at the upper canopy (2 ft from the light source). Arrows indicate the peak enriched wavelengths relative to the control spectrum. **(B)** Representative images of *C. sativa* plants cultivated in growth tents under each light spectrum. **(C)** Morphology of top inflorescences of Pineapple Cough (PC) and Rocky Fire #7 (RF) during the late flowering stage (Week 10). Representative images are shown. Scale bar = 3 cm.

### 2.2 Plant growth and morphology

Plant height was measured from the top of the soil surface to the apex of the top-most inflorescence with a tape measure, and the stem width was measured 5 cm above the surface of the soil using a digital caliper. Representative plant images were taken with a digital camera (EOS Rebel XSi, Canon). At the termination of the experiment, the above-ground plant tissue was harvested and separated into fan leaves, stems, and inflorescences. Fan leaves were dried at 65°C for 5 days (Incubator Owen 310, Robbins Scientific). Stems and inflorescences were hung upside down, and air dried for 14 days in a dark environment at 20°C and 50% humidity following conventional commercial practices. In each case, node number and internode distance were quantified on the main stem. The total dry biomass and that of each tissue separately was measured using an analytical balance (ME104, Mettler Toledo).

### 2.3 Pigment extraction

Photosynthetic pigments (chlorophyll *a*, chlorophyll *b*, carotenoids) were extracted from fully developed fan leaves during the late flowering phase (Week 10). Two leaves of comparable size located directly below the top inflorescence were collected from each plant and stored under liquid N_2_. Frozen leaf tissue was ground, and 25 mg of fine powder was suspended in 1 mL of 90% (v/v) acetone. The solution was thoroughly vortexed, incubated at 4°C for 15 minutes in the dark, and centrifuged (3,000 rcf, 4°C, 5 min) to remove cellular material. The supernatant was diluted 10x and the absorbance was measured spectroscopically at 467, 664 and 647 nm (Cary 60, Agilent Technologies). Chlorophyll (Chl) *a* and *b* content was calculated according to Jeffrey and Humphrey (1975). Carotenoid (Car) content was calculated according to Sun et al., (1998), with the absorbance maximum shifted from ß-carotene to astaxanthin.

### 2.4 Trichome density and morphology

To quantify the density of glandular trichomes, three representative sugar leaves from each plant were collected and carefully placed adaxial side up on black felt to increase the contrast during imaging. Each bract was imaged under UV light (NIGHTSEA, Electron Microscopy Sciences) in a dark room, as previously described (Sutton et al., 2023). Images were acquired using a tripod-mounted digital camera (D3300, Nikon) with a 18-55 mm AF-S DX VR Nikkor lens (Nikon) and two extension tubes (12 mm, 36 mm, JJC Equipment Co.) at 1/20 s exposure, high ISO, f/14, and by using a wireless trigger to minimize image distortion. iLastik (Berg et al., 2019) was used to train a pixel classifier with three classes (leaf, trichomes, background) and to generate probability maps of each class for each image. The leaf area and the trichome count was obtained with Analyze Particles in Fiji (Schindelin et al., 2012) using the probability maps representing “leaf” and “trichomes”.

Scanning electron microscope (SEM) images were used for analyses of glandular trichome morphology. In all cases, nine representative sugar leaves (0.75-1.75 cm) per plant were collected from the terminal inflorescences. The tissue was fixed with 2% glutaraldehyde (v/v) in 0.05 M phosphate buffer (pH 6.8), treated with vacuum to remove gas bubbles, and stored at 4°C for 24 hours. Samples were washed three times with 0.05 M phosphate buffer and dehydrated in increasing concentrations of ethanol (v/v) for three minutes each: 30%, 50%, 70%, 80%, 90%, 95%, 99%, and 3× 100%. Samples were dried with solvent-substituted liquid CO_2_ (Autosamdri 815B critical point dryer, Tousimis), mounted on aluminum specimen stubs with carbon tape, and coated with 4 nm of platinum using a high-resolution sputter coater (208carbon, Cressington). Three sugar leaves were mounted on each stub and imaged (Crossbeam 350 Scanning Electron Microscope, Zeiss) at 2 kV with high current, at a working distance of ∼5.1mm, using mainly the SE2 image signal with some mixing of inlens and SE2 for increased sharpening in the focal plane. Trichome stalk length and glandular head diameter we measured in Fiji (Schindelin et al., 2012). Stalk length was defined as the distance from the epidermal surface to the basal face of the glandular hear, and head diameter was determined as the maximum width of the secretory head.

### 2.5 Cannabinoid extraction and quantification

To measure cannabinoid content, six inflorescence samples per plant from the upper, middle, and lower canopy were sampled and pooled. Upper canopy samples were obtained from the apex inflorescence on the main stem, and the first inflorescence immediately below. Mid-canopy samples were taken from apical inflorescences on branches approximately halfway up the main stem, and lower canopy samples were obtained from apical inflorescences on the lowest branches. Pooled samples were air dried for 14 days in the dark at 20°C and 50% humidity following conventional commercial practices, and ground into fine powder under liquid N_2_. Ground tissue (0.5 g) was extracted twice with 25 mL of 99% EtOH in an ultrasonic bath for 10 minutes, and centrifuged (10 min, 3,900 rcf) to remove tissue debris. Supernatants from both extractions were combined, adjusted to 50 mL with 99% EtOH in volumetric flasks. Samples (1 ml) were then filtered with 0.22 µm PTFE filters into glass HPLC vials and stored at 4°C until analysis.

High-performance liquid chromatography (HPLC) analysis was performed using a Prominence-i LC-2030C-plus HPLC-UV system (Shimadzu), coupled with a Kinetex Polar C18 column (150 x 2.1 mm, 2.6 μm particle size) and a Polar C18 Guard column (Phenomenex Canada). Chromatographic separation was achieved using a three solvent mobile phase of water, methanol and acetonitrile (H_2_O:MeOH:ACN). The gradient was initiated at 36% H_2_O, 36% MeOH and 28% ACN (36:36:28), gradually changed to 27:27:46 over 16 minutes, and then gradually changed to 100% acetonitrile over 8 minutes, in the presence of 0.1% (v/v) trifluoroacetic acid. The column was maintained at 37°C with a flow rate of 0.4 mL/min, with UV detection at 225 nm. The column was washed with 100% acetonitrile for 3 minutes and returned to initial conditions for a 4-minute re-equilibration between samples. To generate calibration curves, cannabinoid standards (CBD, CBG, CBDA, CBGA, CBN, THCVA, Δ9-THC, CBNA, THCA, and CBCA; Cayman Chemical) were injected at a range of concentrations expected for industry-relevant cannabis samples (THCA = 10, 40, 100, 200, 400, 1000 μg/ml, all other cannabinoids = 2, 10, 40, 100, 200, 400 μg/ml). Quantification was performed using peak area integration using LabSolutions (Shimadzu). Cannabinoid content was expressed as concentration per unit inflorescence dry weight (mg/g). Total cannabinoid yield was calculated as cannabinoid concentration per total inflorescence dry weight per plant.

### 2.6 Statistical analyses

All statistical analyses were conducted in R (R Core Team, 2025). Data were organized by tidyverse packages (Wickham et al., 2019). Linear models were fitted with the lm function, with model assumptions and fit confirmed via the plot function (Figure S1) and summarized by the modelsummary::msummary function (Arel-Bundock, 2022). Estimated marginal means and 95% confidence intervals were calculated via the emmeans package (Lenth, 2025). 95% prediction intervals were calculated for each linear model with the predict function to display the variance of estimated individuals from any population. Estimated marginal means and the associated intervals on the real scale were calculated by holding the mean change in height constant across treatments, to reduce interindividual variation. Parameters associated with plant growth and morphology, leaf pigment content, and cannabinoid content were analyzed by one-way ANOVA and Dunnett’s post-hoc test using anova and DescTools::DunnettTest (Signorell, 2025) for values on the real scale, and Kruskal-Wallis test and Dunn’s post-hoc test using kruskal.test and DescTools::DunnTest on the relative scale due to not meeting the assumptions of parametric tests. Plots were constructed with ggplot2 (Wickham, 2016) and ggpubr (Kassambara, 2025).

## 3. Results

In this work we examined the effects of blue and far-red light supplementation on two high-THC *C. sativa* varieties. To achieve this, a typical white light LED spectrum was enriched with blue light (maximum peak at 450 nm) and far-red light (maximum peak at 730 nm) (Figure 1A, 1B). This generated lower R:B ratios in the blue enriched treatment (1.8) compared to the control (2.5) and lower R:FR ratios in the far-red enriched treatment (2.3) compared to the control (10.8) (Table S1). We standardized the cultivation light intensity at canopy level to ensure that spectral quality rather than intensity accounted for any morphological and biochemical variation.

### 3.1 Light spectrum has a limited effect on morphology but regulates pigmentation

First, we observed the morphological variation in the two genotypes, Pineapple Cough and Rocky Fire #7, cultivated under control broad spectrum light. On average, Pineapple Cough plants were taller (∼86 cm), with thinner stems (∼13 cm) and displayed wider apical inflorescences with smaller sugar leaves within (Figure 1C, Table 1). In comparison, Rocky Fire plants were shorter (∼81 cm), with thicker stems (∼16 cm) and with slimmer inflorescences and larger leaves (Figure 1C, Table 1). Cultivation under blue or far-red enriched spectra did not have a significant effect on the absolute value for stem height or width, compared to plants grown under broad spectrum light. Blue light enrichment led to the development of more nodes with shorter internode distances, but the effect was significant only in Pineapple Cough (Table 1). Conversely, the Pineapple Cough (but again, not Rocky Fire) cultivated under far-red enrichment displayed longer internode distances but developed similar number of nodes compared to plants cultivated under standard white spectrum (Table 1).

**Table 1:** Quantification of the morphological characteristics in two *C. sativa* strains cultivated under different light treatments. All measurements were taken at harvest. Data are means ±SD of at least 6 individual plants. Statistically significant differences in the parameters are measured within each strain and compared to the control (white light treatment) were determined with a one-way ANOVA with Dunnett’s post-hoc test (p<0.05) and are bolded

| Treatment | Height (cm) | Stem width (mm) | Node distance (mm) | Nodes (#) |
| --- | --- | --- | --- | --- |
| Pineapple Cough |  |  |  |  |
| White | 85.57 ± 6.94 | 13.08 ± 0.62 | 3.81 ± 0.35 | 19.5 ± 1.6 |
| Blue enriched | 86.97 ± 10.91 | 14.98 ± 2.30 | **3.25 ± 0.33** | **24.0 ± 4.0** |
| Far-red enriched | 91.29 ± 12.05 | 11.82 ± 0.53 | **4.44 ± 0.21** | 18.2 ± 2.1 |
| Rocky Fire #7 |  |  |  |  |
| White | 80.92 ± 11.75 | 15.70 ± 2.14 | 3.73 ± 0.64 | 19.5 ± 1.4 |
| Blue enriched | 82.96 ± 13.93 | 17.90 ± 1.03 | 4.03 ± 0.69 | 19.8 ± 5.2 |
| Far-red enriched | 84.91 ± 15.36 | 13.98 ± 1.90 | 3.87 ± 0.52 | 18.7 ± 2.3 |

Light enrichment had only a minor, genotype-specific influence on plant morphology, but we observed a significant effect on pigment accumulation in both varieties. In all cases, plants accumulated a significantly less Chl *a*, Chl *b*, and carotenoids when cultivated under far-red enriched light resulting in paler leaf coloration, compared to plants cultivated under white light (Figure 2). Cultivation under blue enriched light resulted in higher pigment concentrations and visually darker leaves (Figure 2A); however, the effect was not statistically significant compared to plants cultivated under white light (Figure 2B).

**Figure 2:**
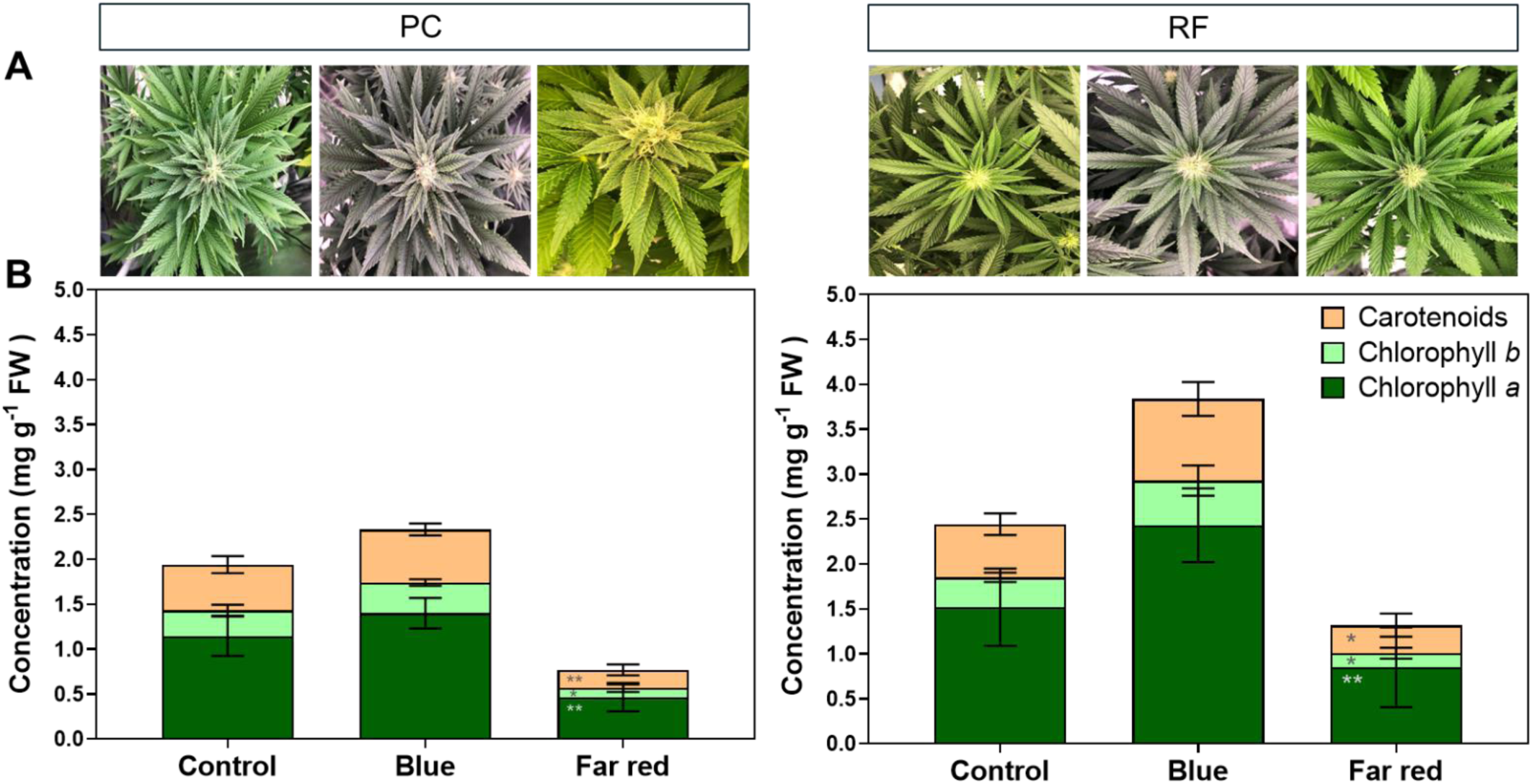
Light spectra influence leaf pigment concentrations. (**A**) Top view of the apical inflorescence of Pineapple Cough (PC) and Rocky Fire #7 (RF) during early flowering (Week 6). (**B**) Concentrations of chlorophyll *a*, chlorophyll *b*, and carotenoids in mature fan leaves collected from the upper canopy at harvest. Data are means ±SD of at least 6 individual plants. Statistically significant differences compared to the control (white light treatment) were determined with a one-way ANOVA with Dunnett’s post-hoc test: * p<0.05; **p<0.01.

### 3.2 Cannabinoid profiles, inflorescence biomass, and total yield

Dry inflorescences were collected from the top, middle and lower canopy and pooled to measure the concentration of major cannabinoids across the plant. The total concentration of cannabinoids per gram of dry inflorescence tissue was on average higher in Pineapple Cough (195 mg g^-1^ DW) when compared to Rocky Fire (145 mg g^-1^ DW) (Figure 3E). Both varieties used in this work are high-THC genotypes, and most of the measured cannabinoids were present as the acidic form of tetrahydrocannabinol (THCA; ∼88% in Pineapple Cough, ∼85% in Rocky Fire), with minor quantities of its decarboxylated form (THC; ∼2% in both strains) and the C3 (varin-type) analog of THCA (THCVA; ∼1.3% in both strains) (Table 2). We also detected small quantities of the cannabinoid precursor CBGA (∼7% in Pineapple Cough, ∼10% in Rocky Fire), and negligible quantities of all other major cannabinoids and their carboxylated forms (CBDA, CBCA, CBNA, CBG). The total concentration of cannabinoids per gram of dry inflorescence tissue was on average higher in Pineapple Cough when compared to Rocky Fire, regardless of the light treatment (Table 2). We also measured the total above-ground plant biomass and tissue allocation between inflorescences, fan leaves, and stems. In all cases, inflorescences accounted for >50% of the total above-ground biomass (Table S2). On average, we observed a higher allocation of biomass towards inflorescences in Pineapple Cough (∼66%) compared to Rocky Fire (∼55%), but total plant biomass did not differ significantly across genotypes or light spectra (Table S2)

**Figure 3.**
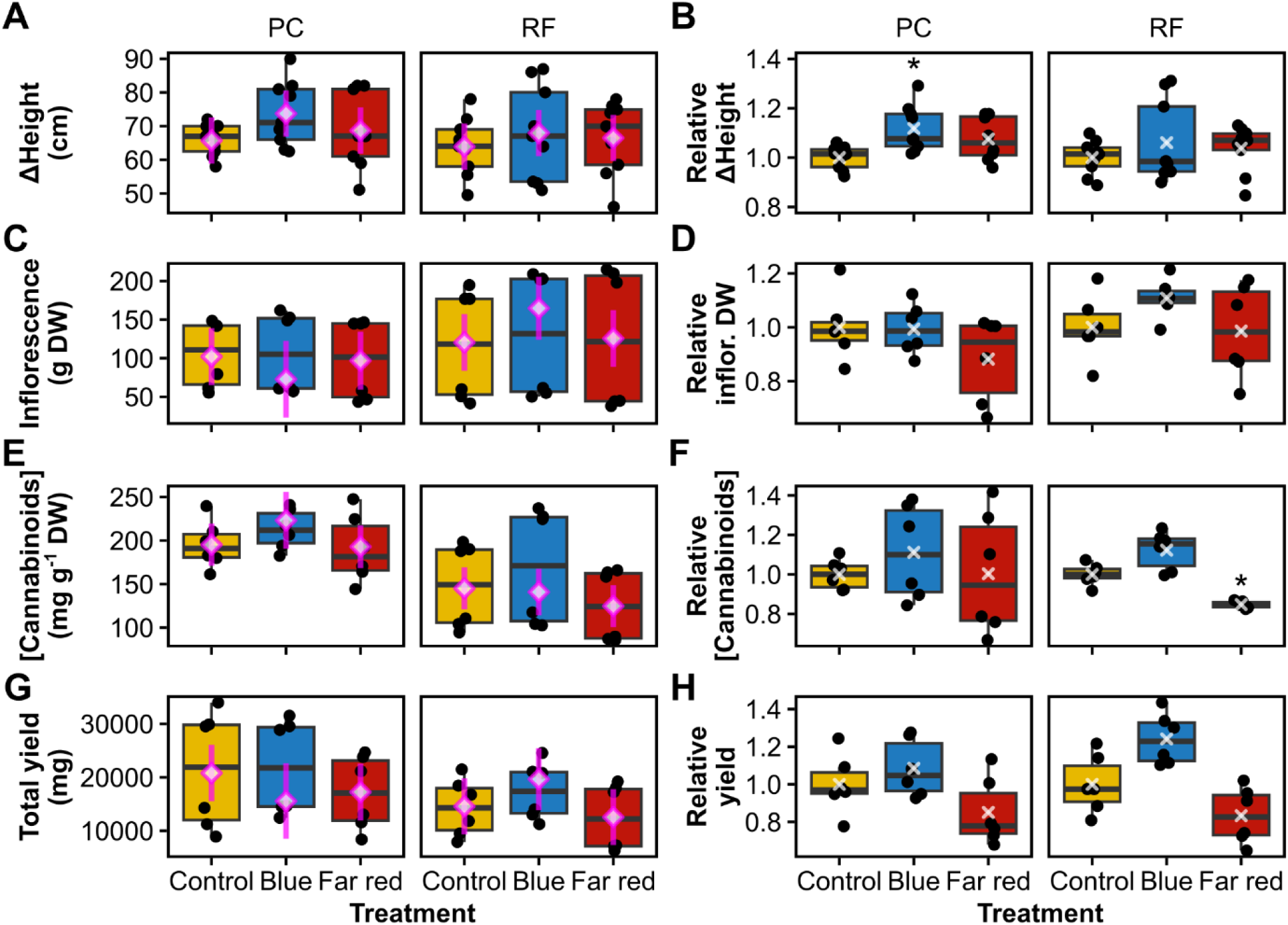
The effect of light spectra on *C. sativa* gross yield metrics on an absolute scale (A,C,E,G) and relative to control plants cultivated under white light (B,D,F,H). **(A)** Change in plant height over 12 weeks of growth. **(B)** Change in height relative to the control treatment. **(C)** Total dry weight of inflorescence tissue per plant. **(D)** Total dry weight of inflorescence tissue per plant relative to the control treatment. **(E)** Concentration of total cannabinoids per inflorescence dry weight. **(F)** Concentration of total cannabinoids per inflorescence dry weigh relative to the control treatment. **(G)** Total cannabinoid yield per plant. **(H)** Total cannabinoids yield per plant relative to the control treatment. White diamonds and magenta lines represent estimated marginal means and 95% confidence intervals, calculated by holding the mean change in height across all groups constant. White X’s represent means. Asterisks denote statistically significant differences from control treatments, calculated with the Kruskal-Wallis test and Dunn’s post hoc test: \**p* < 0.05.

**Table 2:** The concentration of major cannabinoids in inflorescences of two *C. sativa* strains cultivated under different light treatments. In all cases, the numbers represent the cannabinoid concentration per gram of dry inflorescence tissue (mg g^-1^ DW). The data are means ±SD of at least 6 individual plants. Statistical significance was tested with a one-way ANOVA with Dunnett’s post-hoc test, and no differences were found within each strain compared to the control (white light treatment)

| Treatment | THCA | CBDA | CBCA | CBGA | THCVA | THC | CBG | CBNA |
| --- | --- | --- | --- | --- | --- | --- | --- | --- |
| Pineapple Cough |  |  |  |  |  |  |  |  |
| White | 175.3±32.6 | 0.2±0.2 | 2.7±0.7 | 13.4±1.6 | 2.1±2.3 | 4.5±0.8 | 0.8±0.9 | 0.1±0.1 |
| Blue enriched | 190.8±14.1 | 0.3±0.3 | 2.0±0.1 | 15.3±2.9 | 0.5±0.6 | 5.1±3.4 | 0.8±0.9 | 0.1±0.1 |
| Far-red enriched | 171.1±26.8 | 0.2±0.3 | 2.0±0.2 | 12.9±6.9 | 1.1±1.3 | 4.7±2.5 | 0.8±0.9 | 0.1±0.1 |
| Rocky Fire #7 |  |  |  |  |  |  |  |  |
| White | 126.7±43.3 | 0.4±0.2 | 1.0±1.1 | 12.7±0.9 | 2.6±1.2 | 2.9±3.3 | 1.1±0.7 | 0.1±0.2 |
| Blue enriched | 144.4±58.6 | 0.4±0.2 | 1.1±1.2 | 16.4±2.8 | 1.6±0.5 | 3.8±4.2 | 1.2±0.6 | 0.1±0.1 |
| Far-red enriched | 107.0±34.3 | 0.3±0.1 | 1.0±1.1 | 10.6±1.7 | 2.8±1.1 | 2.3±2.6 | 1.0±0.6 | 0.1±0.1 |

Next, we examined the correlation between blue and far-red light enrichment with plant height, floral biomass, cannabinoid concentration, and total yield. The overall plant height after 12 weeks cultivation did not significantly differ between plants grown under different light treatments (Table 1); however, when the starting height of the plants was considered, the change in plant height over the entire cultivation period differed in a genotype-dependent manner (Table S3). We observed a general trend towards greater change in height in Pineapple Cough plants cultivated under blue enriched light compared to control plants cultivated under white light. These trends were weaker and not statistically significant in Rocky Fire (Figure 3A, 3B, Table S3, S4). For inflorescence biomass, cultivation under blue-enriched spectra led to an increase for Rocky Fire on the absolute scale (Figure 3C, Table S3), while that of Pineapple Cough was not significantly different than that measured in control plants (Figure 3C, 3D, Table S3).

We observed that only blue-enriched light led to an increase in cannabinoid concentration in Pineapple Cough compared to the control (Figure 3E, Table S3). In addition, both varieties accumulated less cannabinoids when cultivated under a far-red-enriched spectrum, but this difference was only significant in Rocky Fire relative to control plants (Figure 3F). To estimate the total cannabinoid yield, we multiplied the inflorescence dry weight by the cannabinoid concentration but found no statistically significant differences under various light treatments (Figure 3G, 3H, Table S3, S4). Because inflorescence biomass and total cannabinoid yield did not respond consistently to light treatment, we grouped data from both strains and created more linear models with the change in plant height over time (Δheight; cm) and light treatment as explanatory variables (Table S5, S6). These calculations revealed that the inflorescence biomass and total cannabinoid yield were positively correlated with the change in plant height, regardless of the light spectra (Figure S2). These linear models predicted that for every cm of growth from the beginning of the vegetative phase, plants can gain an additional 5.8 g of inflorescence dry weight as well as 556 mg of total cannabinoid yield at harvest (Table S5). In summary, it appears that light spectra influence plant height and cannabinoid concentration in a genotype-specific manner, but the final cannabinoid yield responds more strongly to plant growth rather than light treatments.

### 3.3 Light spectrum influences trichome morphology

Most cannabinoids in *C. sativa* are synthesized in the glandular trichomes of female inflorescences (Tanney et al., 2021; Xie et al., 2023). To further examine the relationship between light spectrum and cannabis productivity, we observed the density and morphology of the glandular trichomes on sugar leaves. Mature stalked glandular trichomes exhibit intrinsic blue autofluorescence (∼460 nm) when observed under UV light (Hazekamp et al., 2005; Livingston et al., 2020), and we used this property to quantify the trichome density. Consistent with our cannabinoid analysis (Table 2, Figure 3), we observed only minor, non-significant differences in trichome density in plants cultivated under blue and far-red light compared to their control counterparts (Figure 4B). In the autofluorescence imaging, trichomes on plants cultivated under far-red enriched light appeared to have larger heads and longer stalks (Figure 4A), which prompted a detailed investigation of these properties using scanning electron microscopy (SEM).

**Figure 4.**
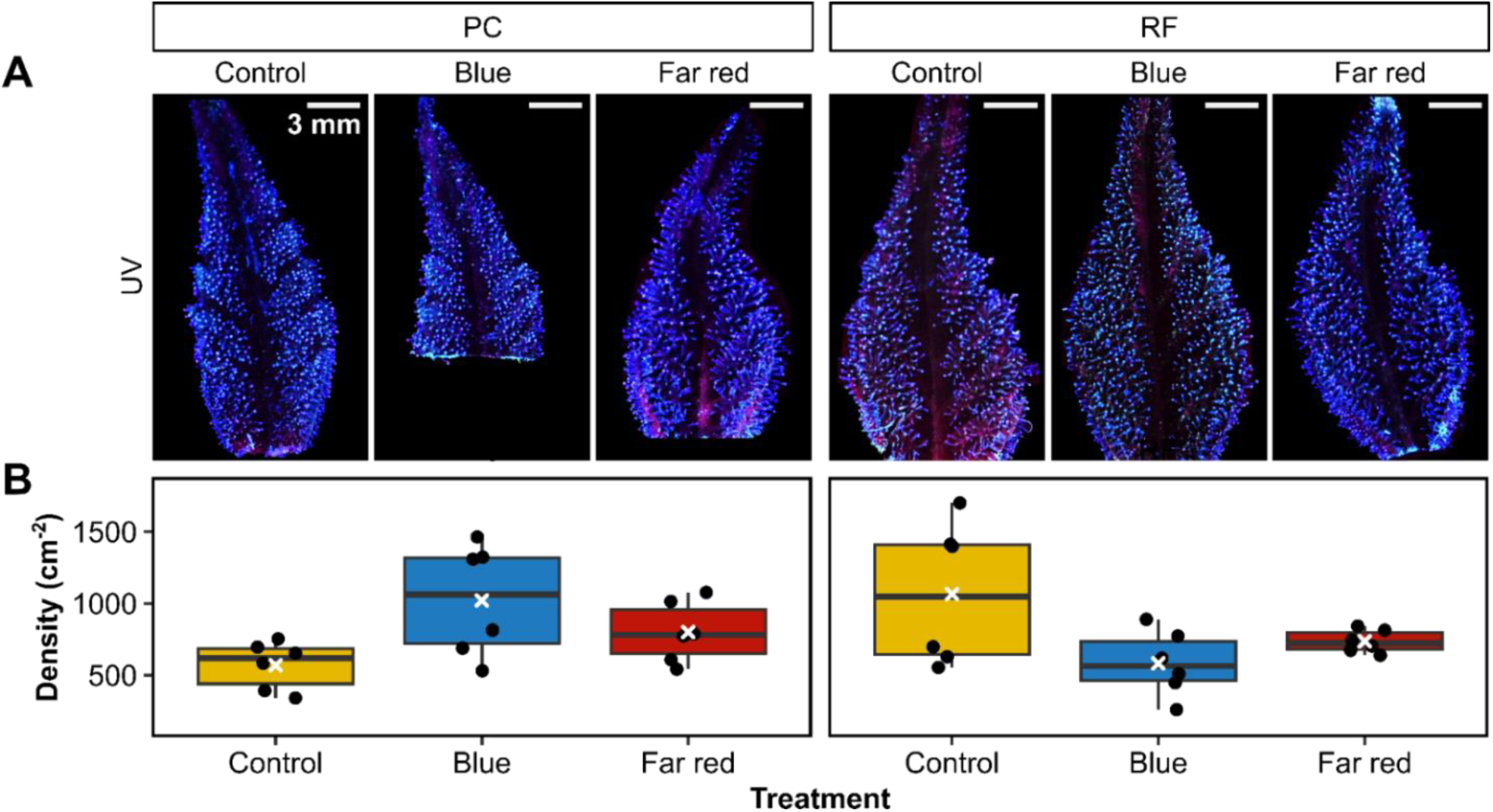
Density of glandular trichomes on sugar leaves. **(A)** Sugar leaves from the apical inflorescence of Pineapple Cough (PC) and Rocky Fire #7 (RF) grown under different light spectra and imaged under UV illumination (scale bars = 3 mm). **(B)** Trichome density (per cm^2^ bract). White crosses (X) represent mean values

SEM imaging revealed the microscopic details of the stalked glandular trichomes. As previously noted (Livingston et al., 2023), we observed that the trichome heads were broken due to the sample preparation and chemical fixation (Figure 5A). Nevertheless, SEM images allowed for the precise quantification of the morphological details of the trichomes, and we observed a significantly increased trichome stalk length in both varieties cultivated under far-red enriched light compared to plants cultivated under white light (Figure 5A, 5B). Cultivation under blue-enriched light leads to longer trichome stalks in Rocky Fire only (Figure 5B). On the other hand, light-spectra influenced the trichome head diameter in Pineapple Cough only, where blue-enriched light led to small but significant decrease in head diameter while far-red-enriched light resulted in an increased head diameter compared to white light- grown controls (Figure 5C).

**Figure 5.**
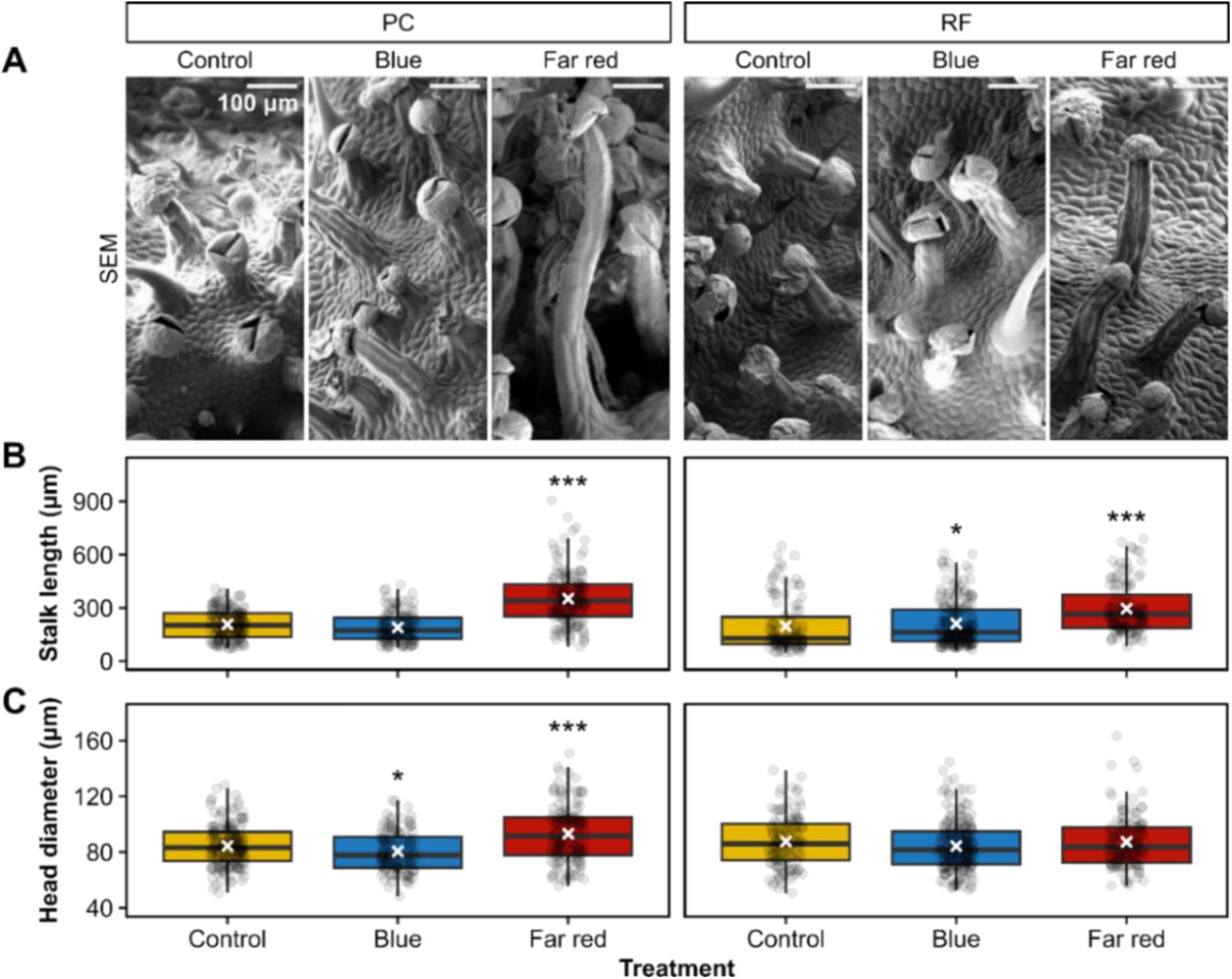
Morphology of glandular trichomes in *C. sativa* cultivated under different light spectra. (**A**) SEM images of glandular trichomes from the apical inflorescence of Pineapple Cough (PC) and Rocky Fire #7 (RF) were imaged with scanning electron microscopy (scale bars = 100 μm). **(B)** Length of glandular trichome stalks and **(C)** trichome head diameter quantified from SEM images. White crosses represent means. Asterisks denote statistically significant differences from control treatments, calculated with post-hoc Dunn’s test: \**p* < 0.05), \*\*\**p* < 0.001.

## 4. Discussion

Optimization of LED lighting strategies has become a major focus of controlled-environment cannabis cultivation, but the enthusiasm for the potential (Ahsan et al., 2024) has not always been supported by empirical data. Several studies testing spectral variation in cannabis have concluded that, unlike intensity, there was no impact of light quality on commercially relevant cannabinoid yield consistently across cannabis varieties (Eaves et al., 2020; Magagnini et al., 2018; Phillips et al., 2025). Here we used commercially available LEDs with enrichment in blue- and far-red spectra to examine the effects on two commercially relevant high-THC *C. sativa* varieties. Our results demonstrated that light quality had a modest and genotype-specific influence on plant morphology but exerted a stronger influence on trichome architecture. While the role of trichome structure and development on cannabinoid biosynthesis has been extensively studied in cannabis (Nolan et al., 2025; Tanney et al., 2021), this is the first report of the influence of light quality on the morphology of these important structures.

### 4.1 Blue and far-red supplementation exert modest influence on *C. sativa* morphology

We observed several trends in light-influenced plant architecture consistent with established photomorphogenic responses to spectral quality, but the magnitude of the morphological responses in our study remained relatively small. Overall plant height, stem width, and total biomass accumulation were only weakly affected by spectral manipulation in both strains (Table 1, S2), but we observed light-induced differences in plant architecture. Far-red light supplementation promoted longer internodes (Table 1), but the response was genotype-specific (only observed in Pineapple Cough). This growth pattern is consistent with the classical shade-avoidance responses mediated through phytochrome and hormone-mediated signaling (Ballaré and Pierik, 2017; Devlin, 2016). Stem elongation attributed to red or far-red light supplementation has been reported in cannabis, albeit this growth pattern is often minor and strain-dependent (Danziger and Bernstein, 2021; Morello et al., 2022; Perotti et al., 2026; Peterswald et al., 2025). Similarly, here we observed only moderate and non-significant increase in overall plant height due to lower R:FR ratios (Figure 3A, 3B, Table 1). Blue light is generally associated with suppression of stem elongation and promotion of compact growth through cryptochrome-mediated pathways (Folta and Carvalho, 2015; Mehmood et al., 2026), consistent with the shorter internodes and increased node density observed in Pineapple Cough (Figure 1, Table 1). Interestingly, we observed a significantly increased relative height in Pineapple Cough under blue-enriched light compared to plants grown under broad spectrum white light (Figure 3B), which contrasts with several previous reports in cannabis in which blue wavelengths suppress elongation growth (Danziger and Bernstein, 2021). It is possible that, in our work, plants cultivated under relatively high and non-limiting PPFD conditions (∼900 µmol m^-2^ s^-1^) were already near optimal levels for growth and biomass accumulation. Under such conditions, spectral quality may function primarily as a developmental modifier rather than a major determinant of growth and biomass accumulation. Finally, our work focused on supplementation of the control broad spectrum white light (Table S1), which may have led to a moderate response in comparison to plants cultivated under monochromatic light (Morello et al., 2022) or with stronger supplementation (e.g. >25% PPFD, Danziger and Bernstein, 2021).

Despite the modest influence of spectral quality on plant growth, cultivation under blue and far-red enrichment resulted in visible pigmentation differences (Figure 1) and photosynthetic pigment concentrations (Figure 2) in both varieties. Far-red enrichment significantly reduced chlorophyll and carotenoid accumulation, resulting in visibly paler leaves, consistent with similar responses in other species (Lauria et al., 2024; Paradiso and Proietti, 2022). Conversely, plants cultivated under blue-enriched spectra has visibly darker leaves and accumulated higher (albeit statistically non-significant) pigment concentrations, as observed in other species (Fantini and Facella, 2020). The effect of light quality on plant physiology via hormone- or photoreceptor-mediated pathways is well described (Chang et al., 2025; de Melo and Alves, 2025; Paik and Huq, 2019), but there is a paucity of similar studies in cannabis. Compared with the strong effects of total irradiance on yield (Chandra et al., 2008; Moher et al., 2022) spectral effects on photosynthetic performance are often modest and genotype-dependent (Danziger and Bernstein, 2021; Morello et al., 2022; Wei et al., 2021; Westmoreland et al., 2021) suggesting that light quality may influence pigmentation rather than large changes in photosynthetic capacity per se.

### 4.2 Light quality does not influence cannabinoid composition or yield

Both strains used in this work accumulate high levels of THCA (Table 2), but we observed only minor differences in cannabinoid profiles, inflorescence biomass, and total yield that could be attributed to blue and far-red supplementation. Cultivation under blue-enriched light generally resulted in higher cannabinoid concentrations and yields, whereas far-red enrichment tended to reduce cannabinoid concentration; however, these responses were modest and variety-specific (Figure 3, Table 2). Our findings generally agree with previous studies reporting that blue-enriched spectra can increase cannabinoid concentrations (Westmoreland et al., 2021) while far-red supplementation was reported to reduce cannabinoid content or production efficiency in a genotype-dependent or moderate manner (Danziger and Bernstein, 2021; Morello et al., 2022; Peterswald et al., 2025; Westmoreland et al., 2021); however, the magnitude of these responses in our work was considerably smaller than reported in some previous investigations.

In the present study, PPFD was maintained at equivalent levels across treatments (Table S1), allowing spectral effects to be evaluated independently of differences in total photon flux. Under these conditions, cannabinoid concentration appeared comparatively stable under different spectral composition, suggesting that total light availability may play a larger role in determining cannabinoid productivity than moderate changes in wavelength distribution alone. Indeed, several studies have reported a strong correlation between total irradiance and yields (Chandra et al., 2008; Eaves et al., 2020; Moher et al., 2022; Rodriguez-Morrison et al., 2021). Furthermore, cannabis varieties exhibit substantial genetic and phenotypic variability, which results in a highly diverse plant architecture and responses to environmental conditions, including spectral quality, even under identical cultivation conditions (Kovalchuk et al., 2020; McKernan et al., 2020; Sawler et al., 2015; Zhang et al., 2018). Thus, cannabis producers must consider that responses to spectral manipulation may not be universally conserved across varieties, and optimization of lighting strategies may require a genotype-specific approach. Further investigation using multiple strains could determine the subtle effects if light quality on cannabis growth and chemical profile and the effect of the genotype on these traits. Establishing light spectral “recipes” based on genotypes may assist in optimizing indoor cultivation strategies.

### 4.3 Spectral quality alters glandular trichome morphology in *C. sativa*

We observed the largest response to spectral manipulation at the level of glandular trichome morphology. While cannabinoid concentration (Table 2, Figure 3), and trichome density (Figure 4) were not different in control, blue- and far-red-enriched spectra, we detected significant differences in trichome stalk length and glandular head diameter. Far-red enrichment consistently increased stalk length in both cultivars and increased glandular head diameter in Pineapple Cough (Figure 5), suggesting that glandular trichome architecture is responsive to spectral quality. Because cannabinoids are synthesized and stored within the secretory disc cells of the glandular trichomes (Livingston et al., 2020; Punja et al., 2023; Tanney et al., 2021) these results suggest that light quality may influence developmental aspects of trichome formation even when overall cannabinoid accumulation remains relatively unchanged. In other glandular trichome-bearing species such as tomato and *Artemisia annua*, light environment influences not only trichome abundance but also glandular development and specialized metabolite production (Chen et al., 2025; Contreras-Avilés et al., 2024; Han et al., 2022) indicating that secretory structures remain developmentally responsive throughout differentiation rather than being fixed once initiated.

The elongation of trichome stalks observed under far-red enrichment (Figure 5) is particularly intriguing because it resembles the cell-expansion responses associated with shade-avoidance growth elsewhere in the plant. Given the minor effects on trichome density (Figure 4), this finding suggest that spectral quality may influence later stages of trichome development, including stalk elongation and glandular head expansion. Whether these responses arise through altered rates of cell expansion, changes in trichome developmental programs, or broader modifications in epidermal differentiation remains unknown. Previous work on *C. sativa* has focused primarily on trichome density and cannabinoid content (Llewellyn et al., 2022; Punja et al., 2023), and this is the first study to directly examine how light spectra influence trichome morphology in cannabis. Detailed examination of the role of light in trichome development is needed to expand on these novel insights.

## 5. Conclusions

Blue and far-red light supplementation exerted relatively modest and variety-dependent effects on cannabis morphology, pigmentation, cannabinoid composition, and yield under controlled environment cultivation with non-limiting growth light intensity. In contrast, glandular trichome morphology responded strongly to spectral manipulation, with far-red enrichment consistently increasing trichome stalk length and, in some cases, glandular head diameter. These findings suggest that trichome development may be more sensitive to spectral quality than cannabinoid accumulation itself. We identify glandular trichomes as a previously underexplored target of light-mediated regulation in cannabis. The functional significance of increased trichome stalk length remains unclear; however, altered trichome architecture could influence metabolite storage, volatile release, or the efficiency of trichome recovery during post-harvest processing. Determining whether these structural modifications confer agronomic advantages will require future studies. Integrating developmental, molecular, and metabolomic approaches will be necessary to determine the mechanisms underlying these morphological responses and whether spectral manipulation can be used to optimize trichome function and phytocannabinoid production in commercial cultivation.

## CRediT Authorship Contribution Statement

**S. Dlaymi**: Investigation, data curation, analysis, visualization, writing; **R.J. Perovich**: Investigation, data curation, analysis, visualization, writing; **T. Kuo**: Investigation, data curation, analysis, visualization, writing; **R. Liu**: Methodology; **V. Fetterley**: Investigation, data curation, analysis, visualization; **A. Z.-F. Lee**: Investigation; **C.S. Harris**: Methodology, resources, validation, supervision, writing; **M. Todesco**: Methodology, resources, validation, supervision, writing; **A.L. Samuels**: Conceptualization, funding acquisition, supervision, resources, validation, writing; **M. Cvetkovska**: Conceptualization, funding acquisition, supervision, visualization, resources, validation, writing.

## Acknowledgements

The authors acknowledge the financial support of Green Amber and the Mitacs Accelerate program (Grant number IT37521 to ALS and IT375534 to MC), as well as supplemental support from the Natural Sciences and Engineering Research Council of Canada (Discovery Grant RGPIN-2025-05870 to ALS). We are grateful for the support from the Canada Foundation for Innovation (CFI; Project Number 39233) and Natural Sciences and Engineering Research Council of Canada CREATE Program (Quality Assurance and Quality Control for Cannabis; Award No: 543319 to MC). We also acknowledge the contributions of the University of Ottawa Greenhouse Facility for support in plant cultivation and the infrastructure and technical assistance of staff in the UBC Bioimaging Facility (RRID: SCR_021304).

## Data Availability

Data will be made available on request.

## References

Ahrens, A., Llewellyn, D., Zheng, Y., 2023. Is Twelve Hours Really the Optimum Photoperiod for Promoting Flowering in Indoor-Grown Cultivars of Cannabis sativa? Plants 12, 2605. 10.3390/plants12142605

Ahsan, S.M., Injamum-Ul-Hoque, M., Shaffique, S., Ayoobi, A., Rahman, M.A., Rahman, M.M., Choi, H.W., 2024. Illuminating Cannabis sativa L.: The Power of Light in Enhancing *C. sativa* Growth and Secondary Metabolite Production. Plants 13, 2774. 10.3390/plants13192774

Andre, C.M., Hausman, J.-F., Guerriero, G., 2016. *Cannabis sativa*: The Plant of the Thousand and One Molecules. Front. Plant Sci. 7. 10.3389/fpls.2016.00019

Arel-Bundock, V., 2022. modelsummary: Data and Model Summaries in R. Journal of Statistical Software 103, 1–23. 10.18637/jss.v103.i01

Ballaré, C.L., Pierik, R., 2017. The shade-avoidance syndrome: multiple signals and ecological consequences. Plant Cell Environ 40, 2530–2543. 10.1111/pce.12914

Berg, S., Kutra, D., Kroeger, T., Straehle, C.N., Kausler, B.X., Haubold, C., Schiegg, M., Ales, J., Beier, T., Rudy, M., Eren, K., Cervantes, J.I., Xu, B., Beuttenmueller, F., Wolny, A., Zhang, C., Koethe, U., Hamprecht, F.A., Kreshuk, A., 2019. ilastik: interactive machine learning for (bio)image analysis. Nat Methods 16, 1226–1232. 10.1038/s41592-019-0582-9

Chandra, S., Lata, H., Khan, I.A., Elsohly, M.A., 2008. Photosynthetic response of *Cannabis sativa* L. to variations in photosynthetic photon flux densities, temperature and CO_2_ conditions. Physiol Mol Biol Plants 14, 299–306. 10.1007/s12298-008-0027-x

Chang, G., Xiang, F., Fan, Y., Li, J., Zhong, S., 2025. Light signal transduction in plants: insights from phytochrome nuclear translocation and photobody formation. New Phytologist 248, 2221–2235. 10.1111/nph.70572

Chen, T., Ma, Y., Qi, J., 2025. Unraveling the Complexity of Plant Trichomes: Models, Mechanisms, and Bioengineering Strategies. Int J Mol Sci 26, 7008. 10.3390/ijms26147008

Contreras-Avilés, W., Heuvelink, E., Marcelis, L.F.M., Kappers, I.F., 2024. Ménage à trois: light, terpenoids, and quality of plants. Trends in Plant Science 29, 572–588. 10.1016/j.tplants.2024.02.007

Danziger, N., Bernstein, N., 2021. Light matters: Effect of light spectra on cannabinoid profile and plant development of medical cannabis (*Cannabis sativa* L.). Industrial Crops and Products 164, 113351. 10.1016/j.indcrop.2021.113351

de Melo, H.C., Alves, F.R.R., 2025. Plant photoreceptors mediate multimodal environmental signaling. Discov. Plants 2, 356. 10.1007/s44372-025-00446-3

Devlin, P.F., 2016. Plants wait for the lights to change to red. Proc Natl Acad Sci USA 113, 7301–7303. 10.1073/pnas.1608237113

Eaves, J., Eaves, S., Morphy, C., Murray, C., 2020. The relationship between light intensity, cannabis yields, and profitability. Agronomy Journal 112, 1466–1470. 10.1002/agj2.20008

Eichhorn Bilodeau, S., Wu, B.-S., Rufyikiri, A.-S., MacPherson, S., Lefsrud, M., 2019. An Update on Plant Photobiology and Implications for Cannabis Production. Front. Plant Sci. 10, 296. 10.3389/fpls.2019.00296

Fantini, E., Facella, P., 2020. Cryptochromes in the field: how blue light influences crop development. Physiologia Plantarum 169, 336–346. 10.1111/ppl.13088

Folta, K.M., Carvalho, S.D., 2015. Photoreceptors and Control of Horticultural Plant Traits. 10.21273/HORTSCI.50.9.1274

Gülck, T., Møller, B.L., 2020. Phytocannabinoids: Origins and Biosynthesis. Trends in Plant Science 25, 985–1004. 10.1016/j.tplants.2020.05.005

Hall, J., Bhattarai, S.P., Midmore, D.J., 2014. The Effects of Photoperiod on Phenological Development and Yields of Industrial Hemp. Journal of Natural Fibers 11, 87–106. 10.1080/15440478.2013.846840

Han, G., Li, Y., Yang, Z., Wang, C., Zhang, Y., Wang, B., 2022. Molecular Mechanisms of Plant Trichome Development. Front. Plant Sci. 13. 10.3389/fpls.2022.910228

Hawley, D., Graham, T., Stasiak, M., Dixon, M., 2018. Improving Cannabis Bud Quality and Yield with Subcanopy Lighting. 10.21273/HORTSCI13173-18

Hazekamp, A., Peltenburg, A., Verpoorte, R., Giroud, C., 2005. Chromatographic and Spectroscopic Data of Cannabinoids from *Cannabis sativa* L. Journal of Liquid Chromatography & Related Technologies 28, 2361–2382. 10.1080/10826070500187558

Huq, E., Lin, C., Quail, P.H., 2024. Light signaling in plants—a selective history. Plant Physiol 195, 213–231. 10.1093/plphys/kiae110

Hussain, T., Jeena, G., Pitakbut, T., Vasilev, N., Kayser, O., 2021. *Cannabis sativa* research trends, challenges, and new-age perspectives. iScience 24, 103391. 10.1016/j.isci.2021.103391

Jeffrey, S.W., Humphrey, G.F., 1975. New spectrophotometric equations for determining chlorophylls *a*, *b*, *c*1 and *c*2 in higher plants, algae and natural phytoplankton. Biochemie und Physiologie der Pflanzen 167, 191–194. 10.1016/S0015-3796(17)30778-3

Jin, D., Jin, S., Chen, J., 2019. Cannabis Indoor Growing Conditions, Management Practices, and Post-Harvest Treatment: A Review. American Journal of Plant Sciences 10, 925–946. 10.4236/ajps.2019.106067

Kassambara A., 2025. ggpubr: ggplot2 Based Publication Ready Plots. R package version 0.6.1. Available at: https://CRAN.R-project.org/package=ggpubr

Kotiranta, S., Sarka, A., Kotilainen, T., Palonen, P., 2025. Decreasing R:FR ratio in a grow light spectrum increases inflorescence yield but decreases plant specialized metabolite concentrations in *Cannabis sativa*. Environmental and Experimental Botany 229, 106059. 10.1016/j.envexpbot.2024.106059

Kovalchuk, I., Pellino, M., Rigault, P., van Velzen, R., Ebersbach, J., R. Ashnest, J., Mau, M., Schranz, M.E., Alcorn, J., Laprairie, R.B., McKay, J.K., Burbridge, C., Schneider, D., Vergara, D., Kane, N.C., Sharbel, T.F., 2020. The Genomics of *Cannabis* and Its Close Relatives. Annu. Rev. Plant Biol. 71, annurev-arplant-081519-040203. 10.1146/annurev-arplant-081519-040203

Kusuma, P., Westmoreland, F.M., Zhen, S., Bugbee, B., 2021. Photons from NIR LEDs can delay flowering in short-day soybean and *Cannabis*: Implications for phytochrome activity. PLOS ONE 16, e0255232. 10.1371/journal.pone.0255232

Lauria, G., Ceccanti, C., Lo Piccolo, E., El Horri, H., Guidi, L., Lawson, T., Landi, M., 2024. “Metabolight”: how light spectra shape plant growth, development and metabolism. Physiologia Plantarum 176, e14587. 10.1111/ppl.14587

Lenth R.V., 2025. emmeans: Estimated Marginal Means, aka Least-Squares Means. R package version 1.11.2-8. Available at: https://CRAN.R-project.org/package=emmeans

Livingston, S.J., Chou, E.Y., Quilichini, T.D., Page, J.E., Samuels, A.L., 2023. Overcoming the challenges of preserving lipid-rich *Cannabis sativa* L. glandular trichomes for transmission electron microscopy. J Microsc 291, 119–127. 10.1111/jmi.13165

Livingston, S.J., Quilichini, T.D., Booth, J.K., Wong, D.C.J., Rensing, K.H., Laflamme-Yonkman, J., Castellarin, S.D., Bohlmann, J., Page, J.E., Samuels, A.L., 2020. Cannabis glandular trichomes alter morphology and metabolite content during flower maturation. The Plant Journal 101, 37–56. 10.1111/tpj.14516

Llewellyn, D., Golem, S., Foley, E., Dinka, S., Jones, A.M.P., Zheng, Y., 2022. Indoor grown cannabis yield increased proportionally with light intensity, but ultraviolet radiation did not affect yield or cannabinoid content. Front. Plant Sci. 13. 10.3389/fpls.2022.974018

Magagnini, G., Grassi, G., Kotiranta, S., 2018. The Effect of Light Spectrum on the Morphology and Cannabinoid Content of *Cannabis sativa* L. Med Cannabis Cannabinoids 1, 19–27. 10.1159/000489030

Mahlberg, P.G., Kim, E.S., 2004. Accumulation of Cannabinoids in Glandular Trichomes of Cannabis (Cannabaceae). Journal of Industrial Hemp 9, 15–36. 10.1300/J237v09n01_04

McKernan, K.J., Helbert, Y., Kane, L.T., Ebling, H., Zhang, L., Liu, B., Eaton, Z., McLaughlin, S., Kingan, S., Baybayan, P., Concepcion, G., Jordan, M., Riva, A., Barbazuk, W., Harkins, T., 2020. Sequence and annotation of 42 cannabis genomes reveals extensive copy number variation in cannabinoid synthesis and pathogen resistance genes. bioRxiv 2020.01.03.894428. 10.1101/2020.01.03.894428

Mehmood, A., Nazir, H., Luo, Y., Wang, W., Urooj, F., Wu, Y., 2026. Blue Light-Mediated Regulation of Growth Morphogenesis and Phytochemical Pathways in Medicinal Plants. Plant Mol Biol Rep 44, 50. 10.1007/s11105-026-01681-y

Moher, M., Llewellyn, D., Jones, M., Zheng, Y., 2022. Light intensity can be used to modify the growth and morphological characteristics of cannabis during the vegetative stage of indoor production. Industrial Crops and Products 183, 114909. 10.1016/j.indcrop.2022.114909

Morello, V., Brousseau, V.D., Wu, N., Wu, B.-S., MacPherson, S., Lefsrud, M., 2022. Light Quality Impacts Vertical Growth Rate, Phytochemical Yield and Cannabinoid Production Efficiency in *Cannabis sativa*. Plants 11, 2982. 10.3390/plants11212982

Namdar, D., Charuvi, D., Ajjampura, V., Mazuz, M., Ion, A., Kamara, I., Koltai, H., 2019. LED lighting affects the composition and biological activity of *Cannabis sativa* secondary metabolites. Industrial Crops and Products 132, 177–185. 10.1016/j.indcrop.2019.02.016

Nolan, M., Guo, Q., Liu, L., Dimopoulos, N., Garcia-de Heer, L., Barkla, B.J., Kretzschmar, T., 2025. Characterisation of Cannabis glandular trichome development reveals distinct features of cannabinoid biosynthesis. Plant Cell Rep 44, 30. 10.1007/s00299-024-03410-9

Paik, I., Huq, E., 2019. Plant photoreceptors: Multi-functional sensory proteins and their signaling networks. Semin Cell Dev Biol 92, 114–121. 10.1016/j.semcdb.2019.03.007

Paradiso, R., Proietti, S., 2022. Light-Quality Manipulation to Control Plant Growth and Photomorphogenesis in Greenhouse Horticulture: The State of the Art and the Opportunities of Modern LED Systems. J Plant Growth Regul 41, 742–780. 10.1007/s00344-021-10337-y

Perotti, F., Pennisi, G., Landolfo, M., Gravina, C., Menozzi, W., Gianquinto, G., Orsini, F., 2026. Optimizing Light Spectra for Cannabis: Effects of End-of-Day and Continuous Far-Red on Plant Morphology and Flower Induction. Horticulturae 12, 456. 10.3390/horticulturae12040456

Peterswald, T.J., Mieog, J.C., Azman Halimi, R., Magner, N.J., Trebilco, A., Kretzschmar, T., Purdy, S.J., 2023. Moving Away from 12:12; the Effect of Different Photoperiods on Biomass Yield and Cannabinoids in Medicinal Cannabis. Plants 12, 1061. 10.3390/plants12051061

Peterswald, T.J., Mieog, J.C., Kretzschmar, T., Purdy, S.J., 2025. The effects of far-red light on medicinal Cannabis. Sci Rep 15, 17435. 10.1038/s41598-025-99771-6

Phillips, A.L., Gill, A., McGorm, B., Burton, R.A., 2025. LED spectra and defoliation independently shape canopy architecture and cannabinoid yield in indoor *Cannabis* cultivation. Industrial Crops and Products 236, 121918. 10.1016/j.indcrop.2025.121918

Punja, Z.K., Sutton, D.B., Kim, T., 2023. Glandular trichome development, morphology, and maturation are influenced by plant age and genotype in high THC-containing cannabis (*Cannabis sativa* L.) inflorescences. J Cannabis Res 5, 12. 10.1186/s42238-023-00178-9

Rodriguez-Morrison, V., Llewellyn, D., Zheng, Y., 2021. Cannabis Inflorescence Yield and Cannabinoid Concentration Are Not Increased with Exposure to Short-Wavelength Ultraviolet-B Radiation. Front. Plant Sci. 12. 10.3389/fpls.2021.725078

Sawler, J., Stout, J.M., Gardner, K.M., Hudson, D., Vidmar, J., Butler, L., Page, J.E., Myles, S., 2015. The Genetic Structure of Marijuana and Hemp. PLOS ONE 10, e0133292. 10.1371/journal.pone.0133292

Schindelin, J., Arganda-Carreras, I., Frise, E., Kaynig, V., Longair, M., Pietzsch, T., Preibisch, S., Rueden, C., Saalfeld, S., Schmid, B., Tinevez, J.-Y., White, D.J., Hartenstein, V., Eliceiri, K., Tomancak, P., Cardona, A., 2012. Fiji: an open-source platform for biological-image analysis. Nat Methods 9, 676–682. 10.1038/nmeth.2019

Siazon, P.M., Nolan, M., Guo, Q., Perovich, R., Kretzschmar, T., Mieog, J.C., 2026. Heads Up: Transcriptomics Reveal Functional Roles of Cannabis Glandular Trichome Stalks. Plants 15, 1624. 10.3390/plants15111624

Signorell A., 2025. DescTools: Tools for Descriptive Statistics. R package version 0.99.60. Available at: https://CRAN.R-project.org/package=DescTools

Sirikantaramas, S., Taura, F., Tanaka, Y., Ishikawa, Y., Morimoto, S., Shoyama, Y., 2005. Tetrahydrocannabinolic Acid Synthase, the Enzyme Controlling Marijuana Psychoactivity, is Secreted into the Storage Cavity of the Glandular Trichomes. Plant Cell Physiol 46, 1578–1582. 10.1093/pcp/pci166

Sun, Z., Cunningham, F.X., Gantt, E., 1998. Differential expression of two isopentenyl pyrophosphate isomerases and enhanced carotenoid accumulation in a unicellular chlorophyte. Proc Natl Acad Sci U S A 95, 11482–11488. 10.1073/pnas.95.19.11482

Sutton, D.B., Punja, Z.K., Hamarneh, G., 2023. Characterization of trichome phenotypes to assess maturation and flower development in *Cannabis sativa* L. (cannabis) by automatic trichome gland analysis. Smart Agricultural Technology 3, 100111. 10.1016/j.atech.2022.100111

Tanney, C.A.S., Backer, R., Geitmann, A., Smith, D.L., 2021. Cannabis Glandular Trichomes: A Cellular Metabolite Factory. Front. Plant Sci. 12. 10.3389/fpls.2021.721986

Wei, X., Zhao, X., Long, S., Xiao, Q., Guo, Y., Qiu, C., Qiu, H., Wang, Y., 2021. Wavelengths of LED light affect the growth and cannabidiol content in *Cannabis sativa* L. Industrial Crops and Products 165, 113433. 10.1016/j.indcrop.2021.113433

Westmoreland, F.M., Kusuma, P., Bugbee, B., 2021. Cannabis lighting: Decreasing blue photon fraction increases yield but efficacy is more important for cost effective production of cannabinoids. PLOS ONE 16, e0248988. 10.1371/journal.pone.0248988

Wickham H., 2016. ggplot2: Elegant Graphics for Data Analysis. Springer-Verlag, New York.

Wickham, H., Averick, M., Bryan, J., Chang, W., McGowan, L.D., François, R., Grolemund, G., Hayes, A., Henry, L., Hester, J., Kuhn, M., Pedersen, T.L., Miller, E., Bache, S.M., Müller, K., Ooms, J., Robinson, D., Seidel, D.P., Spinu, V., Takahashi, K., Vaughan, D., Wilke, C., Woo, K., Yutani, H., 2019. Welcome to the Tidyverse. Journal of Open Source Software 4, 1686. 10.21105/joss.01686

Wu, W., Chen, L., Liang, R., Huang, S., Li, X., Huang, B., Luo, H., Zhang, M., Wang, X., Zhu, H., 2025. The role of light in regulating plant growth, development and sugar metabolism: a review. Front. Plant Sci. 15. 10.3389/fpls.2024.1507628

Xie, Z., Mi, Y., Kong, L., Gao, M., Chen, Shanshan, Chen, W., Meng, X., Sun, W., Chen, Shilin, Xu, Z., 2023. Cannabis sativa: origin and history, glandular trichome development, and cannabinoid biosynthesis. Hortic Res 10, uhad150. 10.1093/hr/uhad150

Zhang, Q., Chen, X., Guo, H., Trindade, L.M., Salentijn, E.M.J., Guo, R., Guo, M., Xu, Y., Yang, M., 2018. Latitudinal Adaptation and Genetic Insights Into the Origins of *Cannabis sativa* L. Front. Plant Sci. 9. 10.3389/fpls.2018.01876

