## Supplementary Figures and Tables for "Light spectral quality alters glandular trichome architecture more strongly than cannabinoid accumulation in *Cannabis sativa*"

**SUPPLEMENTARY MATERIAL**

**Table S1:** Spectral properties of the light treatments used in this work, as Photon Flux Density (PFD) in the 380-780 nm range taken 2 ft from the light source. Light intensity is provided at the blue (400-500 nm), green (500-600 nm), red (600-700 nm) and far-red (700-780 nm) portion of the spectrum.

| **Treatment** | **PFD (μmol m^-2^ s^-1^)** | | | | | **Spectral ratios** | |
| --- | --- | --- | --- | --- | --- | --- | --- |
|  | Total | Blue (B) | Green (G) | Red (R) | Far-Red (FR) | R:B | R:FR |
| **White** | 908 | 137 | 398 | 341 | 32 | 2.5 | 10.8 |
| **Blue enriched** | 900 | 189 | 365 | 315 | 31 | 1.8 | 10.1 |
| **Far-red enriched** | 1090 | 146 | 422 | 364 | 158 | 2.5 | 2.3 |

**Table S2:** Tissue allocation of the total above-ground biomass in two *C. sativa* strains cultivated under different light treatments. All measurements were taken at harvest. The dry biomass data is mean±SD of at least 6 individual plants, and the allocation to inflorescences, fan leaves, and stems is expressed as percent of the total biomass. Statistical significance was tested with a one-way ANOVA with Dunnett’s post-hoc test, and no differences were found within each strain compared to the control (white light treatment)

| **Treatment** | **Dry biomass (g)** | | | **Inflorescence (%)** | **Fan Leaf (%)** | **Stem (%)** |
| --- | --- | --- | --- | --- | --- | --- |
| Pineapple Cough | | | | | | |
| **White** | 179.4 | ± | 16.9 | 68 | 17 | 15 |
| **Blue enriched** | 199.2 | ± | 38.4 | 64 | 18 | 18 |
| **Far-red enriched** | 177.6 | ± | 32.8 | 63 | 18 | 19 |
| Rocky Fire #7 | | | | | | |
| **White** | 184.5 | ± | 65.7 | 55 | 27 | 19 |
| **Blue enriched** | 174.2 | ± | 79.4 | 54 | 27 | 19 |
| **Far-red enriched** | 190.2 | ± | 67.8 | 58 | 24 | 18 |

**Figure S1**. Diagnostic plots for linear models used in Figures 3A, C, D, F, and S2.

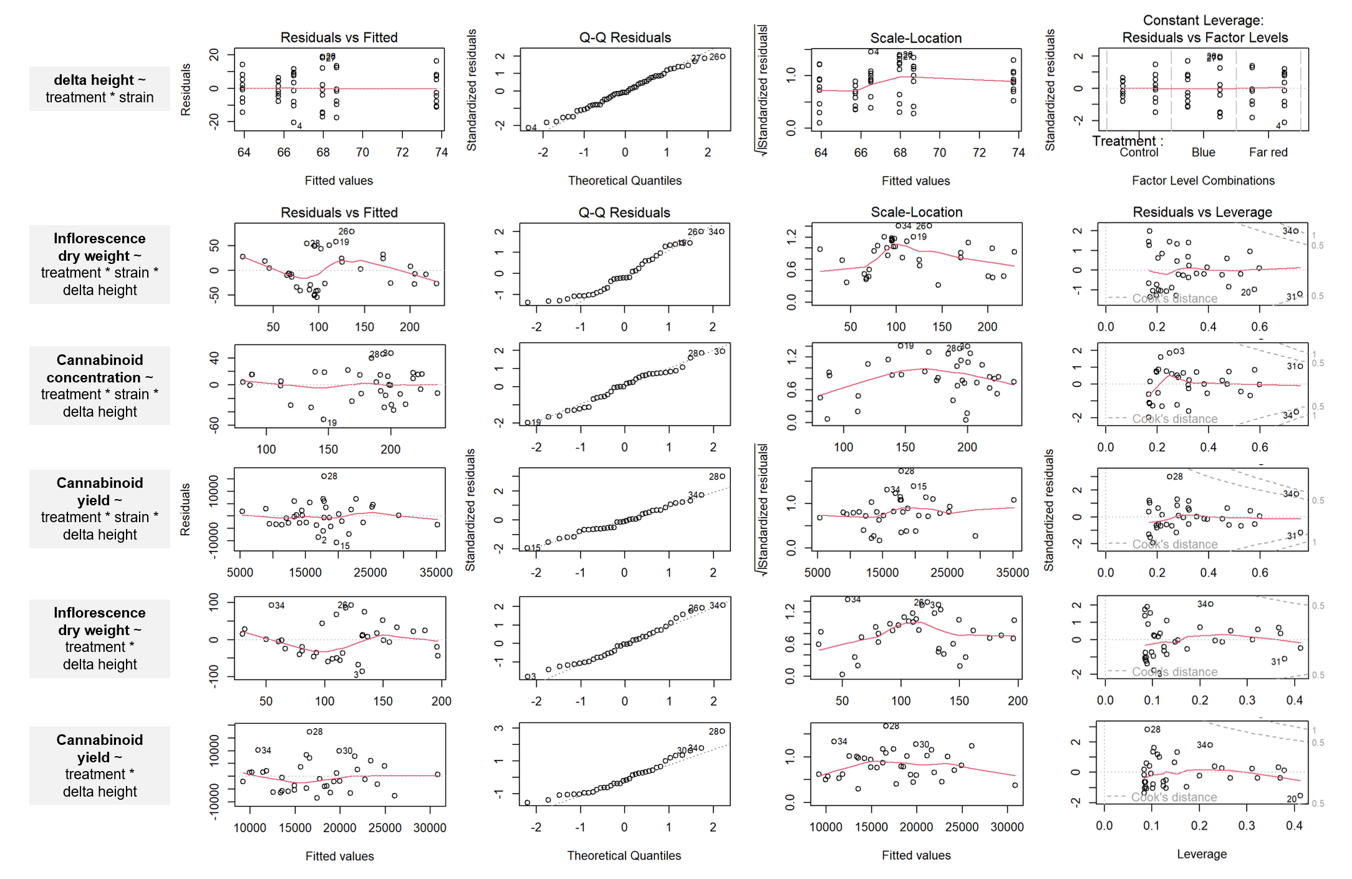

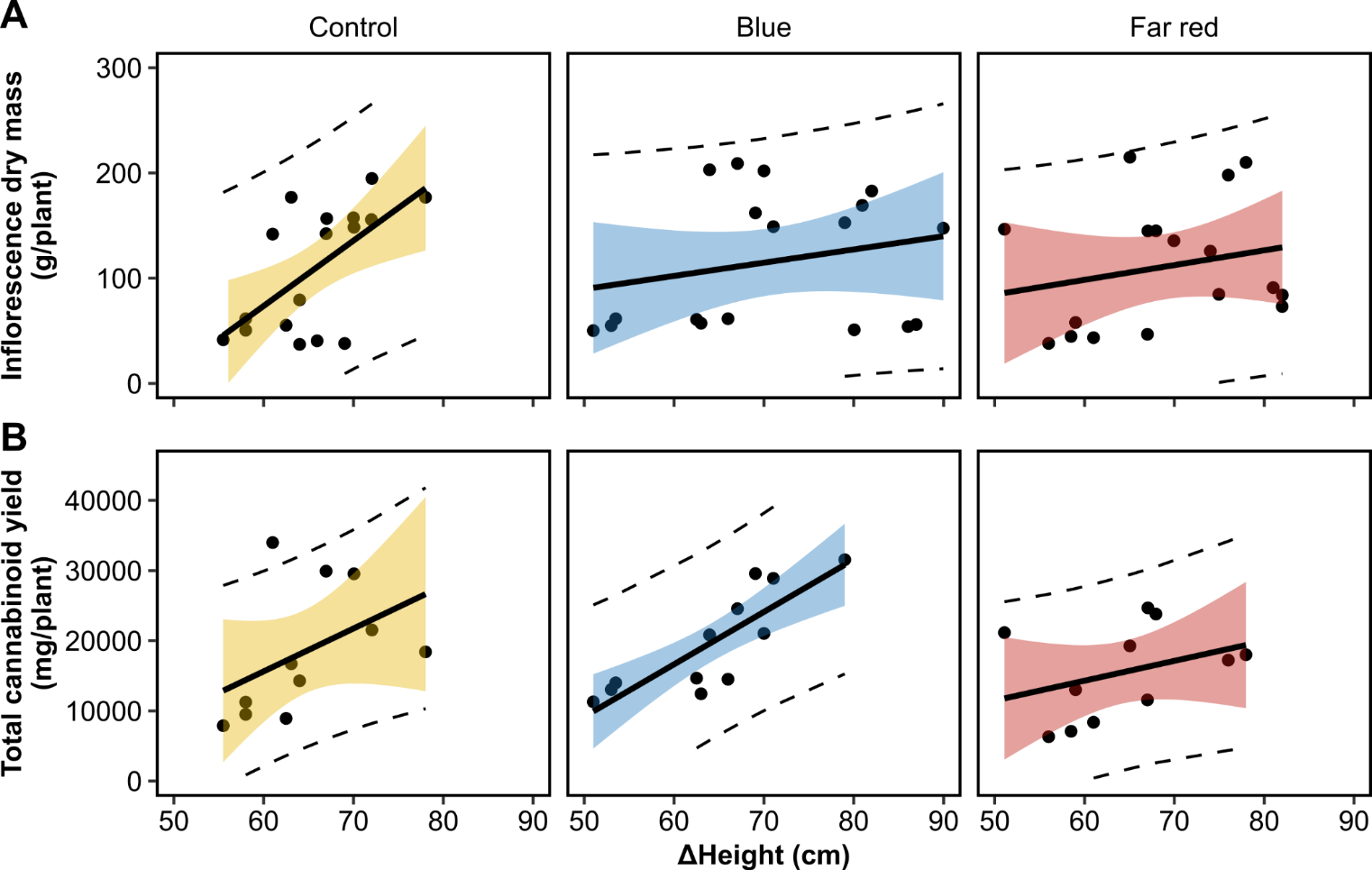
**Figure S2**. Linear model output for **(A)** The inflorescence dry weigh (g) and **(B)** total cannabinoid yield (mg/plant) as response variables, and the change in plant height over time (Δheight; cm) and light spectra as explanatory variables. Colored ribbons represent 95% confidence intervals and dashed lines represent 95% prediction intervals

**Table S3:** A summary of four linear models used in Figures 3A, C, D, G: (1) **Δheight** ~ treatment * strain; (2) **inflorescence dry weight** ~ treatment * strain * Δ height; (3) **cannabinoid concentration** ~ treatment * strain * Δheight; and (4) **cannabinoid yield** ~ treatment * strain * Δheight. Parentheses indicate standard errors.

|  | **Dependent variables** | | | |
| --- | --- | --- | --- | --- |
|  | **Δheight** | **Inflorescence DW** | **Cannabinoid conc.** | **Cannabinoid yield** |
| (Intercept) | 65.724*** | −308.430 | 154.408 | −58985.342 |
|  | (3.414) | (290.412) | (190.890) | (41361.735) |
| TreatmentBlue | 8.000 | −47.496 | 196.879 | −4774.644 |
|  | (4.828) | (362.970) | (238.582) | (51695.654) |
| TreatmentFar red | 2.953 | 433.150 | −86.808 | 68222.282 |
|  | (4.828) | (344.870) | (226.685) | (49117.782) |
| StrainRF | −1.823 | 64.964 | 234.827 | 47779.736 |
|  | (4.828) | (312.709) | (205.546) | (44537.301) |
| TreatmentBlue × StrainRF | −3.951 | −155.533 | 49.505 | −3452.838 |
|  | (6.829) | (407.051) | (267.557) | (57973.862) |
| TreatmentFar red × StrainRF | −0.363 | −462.480 | 10.674 | −70874.629 |
|  | (6.829) | (377.885) | (248.386) | (53819.986) |
| delta_height_cm |  | 6.482 | 0.644 | 1259.639 |
|  |  | (4.547) | (2.989) | (647.589) |
| TreatmentBlue × delta_height_cm |  | 0.286 | −2.667 | −7.416 |
|  |  | (5.543) | (3.644) | (789.518) |
| TreatmentFar red × delta_height_cm |  | −6.921 | 1.335 | −1133.200 |
|  |  | (5.435) | (3.573) | (774.087) |
| StrainRF × delta_height_cm |  | −0.736 | −4.499 | −852.506 |
|  |  | (4.901) | (3.221) | (697.954) |
| TreatmentBlue × StrainRF × delta_height_cm |  | 3.619 | −1.288 | 217.711 |
|  |  | (6.303) | (4.143) | (897.640) |
| TreatmentFar red × StrainRF × delta_height_cm |  | 7.464 | −0.453 | 1143.075 |
|  |  | (5.950) | (3.911) | (847.449) |
| Num.Obs. | 54 | 36 | 36 | 36 |
| R^2^ | 0.093 | 0.681 | 0.779 | 0.567 |
| R^2^ Adj. | −0.002 | 0.534 | 0.677 | 0.369 |
| AIC | 412.2 | 385.5 | 355.3 | 742.5 |
| BIC | 426.1 | 406.0 | 375.8 | 763.1 |
| Log.Lik. | −199.077 | −179.731 | −164.625 | −358.248 |
| F | 0.980 | 4.649 | 7.684 | 2.862 |
| RMSE | 9.66 | 35.64 | 23.43 | 5076.59 |
| *** p < 0.001 |  |  |  |  |

**Table S4:** A summary of estimated marginal means (EMM) displayed in Figures 3A, C, D, G. Within each strain, EMMs that fall outside of the 95% CI of the control group are bolded to indicate significant differences from the control. PC = Pineapple Cough; RF = Rocky Fire #7.

| **Strain** | **Treatment** | **Δheight (cm)** | **EMM** | **SE** | **df** | **95% CI** | |
| --- | --- | --- | --- | --- | --- | --- | --- |
|  |  |  |  |  |  | **Lower** | **Upper** |
| **delta height** ~ treatment * strain | | | | | | | |
| PC | Control |  | 65.7 | 3.4 | 48 | 58.9 | 72.6 |
|  | Blue |  | **73.7** | 3.4 | 48 | 66.9 | 80.6 |
|  | Far red |  | 68.7 | 3.4 | 48 | 61.8 | 75.5 |
| RF | Control |  | 63.9 | 3.4 | 48 | 57.0 | 70.8 |
|  | Blue |  | 68.0 | 3.4 | 48 | 61.1 | 74.8 |
|  | Far red |  | 66.5 | 3.4 | 48 | 59.6 | 73.4 |
| **inflorescence dry weight** ~ treatment * strain * delta height | | | | | | | |
| PC | Control | 63.3 | 102.2 | 17.9 | 24 | 65.2 | 139.2 |
|  | Blue | 63.3 | 72.8 | 24.0 | 24 | 23.1 | 122.4 |
|  | Far red | 63.3 | 96.9 | 18.2 | 24 | 59.4 | 134.4 |
| RF | Control | 63.3 | 120.5 | 17.9 | 24 | 83.6 | 157.4 |
|  | Blue | 63.3 | **164.8** | 19.8 | 24 | 124.0 | 205.6 |
|  | Far red | 63.3 | 125.6 | 17.8 | 24 | 88.8 | 162.3 |
| **cannabinoid concentration** ~ treatment * strain * delta height | | | | | | | |
| PC | Control | 63.3 | 195.2 | 11.8 | 24 | 170.9 | 219.5 |
|  | Blue | 63.3 | **223.2** | 15.8 | 24 | 190.5 | 255.8 |
|  | Far red | 63.3 | 192.9 | 11.9 | 24 | 168.3 | 217.6 |
| RF | Control | 63.3 | 145.1 | 11.7 | 24 | 120.8 | 169.3 |
|  | Blue | 63.3 | 140.9 | 13.0 | 24 | 114.1 | 167.7 |
|  | Far red | 63.3 | 124.8 | 11.7 | 24 | 100.6 | 148.9 |
| **cannabinoid yield** ~ treatment * strain * delta height | | | | | | | |
| PC | Control | 63.3 | 20803.7 | 2552.0 | 24 | 15536.7 | 26070.7 |
|  | Blue | 63.3 | 15559.3 | 3424.7 | 24 | 8491.0 | 22627.5 |
|  | Far red | 63.3 | 17245.9 | 2585.7 | 24 | 11909.2 | 22582.6 |
| RF | Control | 63.3 | 14583.3 | 2544.0 | 24 | 9332.8 | 19833.8 |
|  | Blue | 63.3 | 19676.5 | 2814.8 | 24 | 13867.1 | 25485.8 |
|  | Far red | 63.3 | 12556.4 | 2538.4 | 24 | 7317.5 | 17795.4 |

**Table S5:** A summary of two linear models used in Figure S2: (1) **inflorescence dry weight** ~ treatment * delta height; and (2) **cannabinoid yield** ~ treatment * delta height. Parentheses indicate standard errors.

|  | **Dependent variables** | |
| --- | --- | --- |
|  | **Inflorescence DW** | **Cannabinoid yield** |
| (Intercept) | -253.480 | -17364.159 |
|  | (124.869) | (16005.782) |
| delta_height_cm | 5.762** | 556.489* |
|  | (1.962) | (251.436) |
| TreatmentBlue | 36.747 | -10789.234 |
|  | (172.588) | (22122.457) |
| TreatmentFar red | 59.116 | 11086.255 |
|  | (161.737) | (20731.537) |
| delta_height_cm × TreatmentBlue | -0.529 | 190.088 |
|  | (2.693) | (345.168) |
| delta_height_cm × TreatmentFar red | -0.889 | -220.264 |
|  | (2.546) | (326.292) |
| Num.Obs. | 36 | 36 |
| R^2^ | 0.463 | 0.411 |
| R^2^ Adj. | 0.374 | 0.313 |
| AIC | 392.2 | 741.6 |
| BIC | 403.2 | 752.7 |
| Log.Lik. | -189.078 | -363.802 |
| F | 5.176 | 4.189 |
| RMSE | 46.21 | 5923.37 |
| * p < 0.05, ** p < 0.01 | | |

**Table S6:** A summary of estimated marginal means (EMM) of the data presented in Figure S2.

| **Treatment** | **Delta height (cm)** | **EMM** | **SE** | **df** | **95% CI** | |
| --- | --- | --- | --- | --- | --- | --- |
|  |  |  |  |  | **Lower** | **Upper** |
| **inflorescence dry weight** ~ treatment * delta height | | | | | | |
| Control | 63.3 | 111.5 | 14.6 | 30 | 81.6 | 141.3 |
| Blue | 63.3 | 114.7 | 14.7 | 30 | 84.8 | 144.7 |
| Far red | 63.3 | 114.3 | 14.6 | 30 | 84.3 | 144.2 |
| **cannabinoid yield** ~ treatment * delta height | | | | | | |
| Control | 63.3 | 17885.4 | 1873.4 | 30 | 14059.4 | 21711.4 |
| Blue | 63.3 | 19136.9 | 1881.5 | 30 | 15294.4 | 22979.4 |
| Far red | 63.3 | 15019.5 | 1877.7 | 30 | 11184.9 | 18854.2 |
